# Monitoring of language selection errors in switching: Not all about conflict

**DOI:** 10.1101/357061

**Authors:** Xiaochen Zheng, Ardi Roelofs, Jason Farquhar, Kristin Lemhöfer

**Affiliations:** Donders Institute for Brain, Cognition and Behaviour, Radboud University, Nijmegen, The Netherlands; International Max Planck Research School for Language Sciences, Max Planck Institute for Psycholinguistics, Nijmegen, The Netherlands

**Keywords:** language switching, error monitoring, conflict monitoring, ERN, N2

## Abstract

Although bilingual speakers are very good at selectively using one language rather than another, sometimes language selection errors occur. To investigate how bilinguals monitor their speech errors and control their languages in use, we recorded event-related potentials (ERPs) in unbalanced Dutch-English bilingual speakers in a cued language-switching task. We tested the conflict-based monitoring model by investigating the error-related negativity (ERN) and comparing the effects of the two switching directions (i.e., to the first language, L1 vs. to the second language, L2). Results show that the speakers made more language selection errors when switching from their L2 to the L1 than vice versa. In the EEG, we observed a robust ERN effect following language selection errors compared to correct responses, reflecting monitoring of speech errors. Most interestingly, the ERN effect was enlarged when the speakers were switching to their L2 (less conflict) compared to switching to the L1 (more conflict). Our findings do not support the conflict-based monitoring model. We discuss an alternative account in terms of error prediction and reinforcement learning.

## Introduction

While talking to each other, bilinguals seem to effortlessly switch between languages. Only occasionally does a word from the currently unused language slip into the active language [1, 2], such as a Dutch-English speaker using “misschien”, the Dutch word for “maybe”, when talking to their English-speaking colleagues. The rarity of such slips, or *language selection errors*, suggests strong language control mechanisms which allow bilinguals to separate but also fluently mix two languages if desired [3–5]. As part of the control process, bilinguals also constantly monitor what they have just said and what they are about to say, inspecting speech errors and intervening when necessary [6].

In the context of the monitoring and control process, being able to detect errors is particularly crucial. Investigating such on-line processing mechanisms is possible with the help of event-related potentials (ERPs) in the EEG. Research on action monitoring has reliably shown a negative-going peak elicited by erroneous responses, such as pressing the wrong button in a response selection task [7, 8]. This so-called error-related negativity (ERN) begins around the onset of error commission and peaks around 100 ms thereafter. It has a fronto-central distribution and has been associated with activity in the dorsal anterior cingulate cortex (ACC) or pre-supplementary motor area (pre-SMA), regions which are broadly connected to motor planning and control systems [9, 10].

In the language domain, research has also shown an ERN-like component after a vocal slip [11–13] or after an error in meta-linguistic tasks which require covert production [14]. The first ERN-like component in response to vocal slips was reported by Masaki et al. [12] in a Stroop task. Later, Ganushchak and Schiller [15] reported an ERN following verbal errors in a second language (L2) as well, using a phoneme-monitoring task. Furthermore, Riès and colleagues [13] showed that the ERN component is also present, but smaller, after correct vocal responses (correct response negativity, CRN), suggesting its role in general on-line response monitoring rather than error detection specifically. Nevertheless, relatively few studies on monitoring have been conducted so far with overt production [11–13, 16–18], partially because it is challenging to measure the response-locked EEG when there are motor artifacts caused by articulation (for a review, see [19]). To our knowledge, no EEG studies have been done yet on the monitoring of language selection errors.

The similarity between the findings on the ERN in action monitoring and those in language production monitoring raises the question whether speech monitoring is a special case of domain-general performance monitoring [13, 20–21]. Nozari et al. [21] proposed a conflict-based monitoring mechanism implemented in the interactive two-step model of word production [22], which adapted the conflict monitoring theory of action monitoring [23–24] and applied it to speech production. According to the conflict monitoring theory, error detection is accomplished by monitoring response conflict. The ERN, triggered by such conflicts, functions as a signal for the control system to recruit and regulate the amount of top-down control, in order to resolve conflict and subsequently adapt performance [23–24], During language production, such conflicts can occur during word selection and phoneme selection [21]. According to the conflict-based model, the more conflict arises during word selection or phoneme selection, the more likely one is to make a semantic error (e.g., saying “dog” for “cat”) or a phonological error (e.g., saying “cag” for “cat”), respectively. As a consequence, the ERN as a signal for error/conflict detection will become larger as error rates increase [21] (but see [24]).

The concept of conflict monitoring may also be relevant for bilingual control. During bilingual production, conflicts arise not only within a language (e.g., word selection, phoneme selection), but also between multiple languages which are simultaneously activated [25–28]. Bilingual language control is often investigated using a picture naming task, where bilingual speakers are asked to name pictures in either of their languages according to a language cue (e.g., a flag or a color patch). Usually, bilinguals are faster to name the pictures in their stronger, dominant first language (L1) compared to their weaker L2 [29]. However, in language switching contexts, the language dominance effect is eliminated or sometimes even reversed: When bilingual speakers have to switch between languages, they become slower [30–33] and make more errors [34] when switching to L1 compared to switching to L2. This so-called *reversed dominance effect* is usually explained by inhibition of the nontarget language or/and enhancement of the target language [35]. Such cognitive control is stronger in L2 trials than in L1 trials where the dominant L1 has to be inhibited or/and the weaker L2 has to be enhanced. Therefore, switching back to the L1 requires more effort to overcome the residual control and results in a reversed effect. According to the conflict-based model, the higher reaction times (RTs) and error rates when switching from L2 to L1 also suggest more conflict in this switching direction than the other way around. If speech monitoring is conflict-based, the question arises whether the monitoring process, as reflected in the ERN, will parallel the difference in the amount of conflict between the two switching directions. This question will be addressed by the present study.

## The current study

To investigate how bilinguals monitor their speech errors and control their languages in use, the current study tested the conflict-based monitoring model [21] by examining the ERPs of Dutch-English speakers in a bilingual picture naming task. We were particularly interested in the ERN component as an index of error/conflict detection. According to the conflict-based model, the amount of conflict predicts the probability of error occurrence, and the ERN should increase in high-conflict conditions [21]. Based on previous findings on the reversed dominance effect [31–32, 34], we expect more language selection errors when switching from L2 to L1 than in the opposite switching direction. If the conflict-based model is correct, then the amount of conflict should be higher in that condition, and we should observe a larger ERN following a language selection error in switching from L2 to L1 than vice versa.

Another ERP component that we are interested in is the (stimulus-locked) N2, a negative wave peaking between 200 and 350 ms after stimulus onset. The N2 component is believed to reflect pre-response conflict and shares a similar scalp topography and presumed neural source as the ERN [24]. Therefore, we also expect the N2 to show the same pattern as the ERN in terms of switching directions. It should be noted that the N2 component in language production is also interpreted differently by other researchers as a reflection of inhibitory control [36–37] and overcoming the inhibition during (language) switching [33, 37–38]. In inhibition or overcoming previous inhibition, the N2 has a fronto-central or posterior scalp distribution, respectively (see also [39] for a review on nonlinguistic research).

To further test the conflict-based model, we included another manipulation of response conflict in language production, namely, cognate status. Cognates are words with a form-similar translation equivalent between different languages (e.g., “table” in English and “tafel” in Dutch). Previous bilingual production research has shown slower picture naming and more errors for noncognates than for cognates [29, 40–41], suggesting more conflict in noncognate naming. In addition, according to the model of Nozari et al. [21], there is less conflict when there is form overlap (e.g., there is no conflict between the onset phonemes of “table” and “tafel”), thus the naming of cognates should yield less conflict than that of noncognates. Based on previous literature [41], we expect higher error rates in noncognates compared to cognates. If the conflict-based monitoring model is correct, we should also observe a larger CRN as well as a larger N2 when (correctly) naming noncognates as compared to cognates (but see [42] for the opposite view). To avoid the possible interaction between the cognate effect and the switch effect, we only manipulated cognate status on repeat trials.

## Method

### Participants

Twenty-eight participants took part in the study for course credit or vouchers. All of them were native Dutch speakers, raised monolingually, who spoke English as their most proficient nonnative language. All the participants were right-handed and had normal or corrected-to-normal vision. We excluded the EEG data from four participants either because of excessive artifacts, or because they did not make enough errors for analysis. To be consistent, we also excluded their data from the behavioral analysis. This resulted in a final set of 24 participants (five males). Table 1 shows the language background of the 24 participants as assessed by a questionnaire, and their English vocabulary size measured by the LexTALE test [43].

**Table 1.** Participants’ Language Background and English Proficiency (N = 24).

| Characteristic | *Mean* | *SD* | *Range* |
| --- | --- | --- | --- |
| Age | 22.9 | 2.7 | 19-30 |
| Age of acquiring English | 9.3 | 1.8 | 6-11 |
| Self-rated frequency of using English<sup>a</sup> |  |  |  |
| - speaking | 3.4 | 1.1 | 2-5 |
| - listening | 4.5 | 0.8 | 2-5 |
| - reading | 3.7 | 1.2 | 1-5 |
| - writing | 2.8 | 1.4 | 1-5 |
| Self-rated frequency of switching languages<sup>a</sup> |  |  |  |
| - speaking | 2.0 | 0.9 | 1-4 |
| Self-rated proficiency in English<sup>a</sup> |  |  |  |
| - speaking | 4.3 | 0.7 | 3-5 |
| - listening | 4.5 | 0.6 | 3-5 |
| - writing | 4.1 | 0.9 | 2-5 |
| - reading | 4.5 | 0.6 | 3-5 |
| English vocabulary size |  |  |  |
| - LexTALE test | 77.7 | 10.4 | 62-98 |

*Note.* *SD* = Standard Deviation.
<sup>a</sup>Self-ratings were given on a scale from 1 = *very rare/bad* to 5 = *very often/good*.

### Materials

Experimental stimuli consisted of 40 black-and-white line drawings, representing 20 translation pairs of Dutch-English noncognate words (e.g., Dutch word “boom” and its English translation “tree”) and 20 pairs of cognate words (e.g., Dutch word “tijger” and its English translation “tiger”). We selected the pictures mainly from the international picture naming project (IPNP) database [44], opting for those with highest naming agreement in both Dutch and English [44–45]. The cognate and noncognate words were selected based on Levenshtein distance [46] of phonetic translations and on word etymology. We matched all the Dutch and English, cognate and noncognate picture names as closely as possible on number of syllables (*F* < 1) and the phonological onset categories, so that possible differences in RTs could not be explained by word length or differences in voice-key sensitivity (e.g., /f/ and /s/ have a delayed voice-key onset compared to /p/ and /t/). Within each language, we matched cognate and noncognate words on lemma (log) frequency (both *ps* > .166; CELEX database [47]). Given the restrictions above, we used some additional pictures drawn from scratch. All the pictures were edited to a size of 300 x 300 pixels. Table 2 shows the characteristics for the noncognate and cognate words used in the study. A full list of cognate and noncognate words can be found in S1 Appendix A.

**Table 2.**
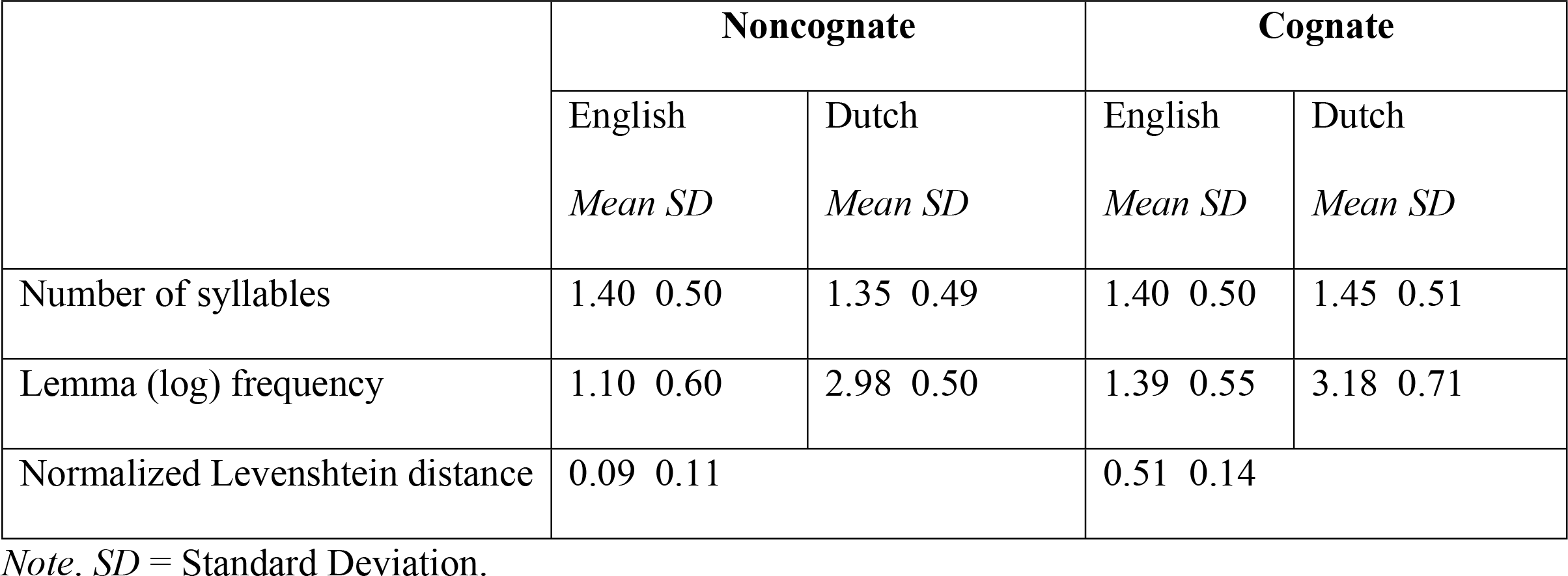
Word Characteristics for Noncognates and Cognates.

|  | Noncognate |  | Cognate |  |
| --- | --- | --- | --- | --- |
|  | English | Dutch | English | Dutch |
|  | *Mean SD* | *Mean SD* | *Mean SD* | *Mean SD* |
| Number of syllables | 1.40 0.50 | 1.35 0.49 | 1.40 0.50 | 1.45 0.51 |
| Lemma (log) frequency | 1.10 0.60 | 2.98 0.50 | 1.39 0.55 | 3.18 0.71 |
| Normalized Levenshtein distance | 0.09 0.11 |  | 0.51 0.14 |  |

*Note.* *SD* = Standard Deviation.

### Design

There were two types of trials: switch trials, where the response language was different from that in the previous trial, and repeat trials, where the response language stayed the same. Depending on which language was required on the current trial, we further categorized switch trials as “switch to Dutch (L1)” and “switch to English (L2)”, and repeat trials as “repeat in Dutch (L1)” and “repeat in English (L2)”. In the current study, we focused on two analyses: on switch trials (that contained noncognates only), we compared language selection errors with correct responses; on repeat trials, we compared correct cognate naming with noncognate naming. We only used a subset of repeat and switch trials for analyses (i.e., critical repeat and critical switch trials). The selection of critical trials is explained below.

Each experimental list contained 640 trials, divided into eight blocks of 80 trials. Each stimulus appeared twice in a block, once in Dutch and once in English. Each list had 160 switch trials (switch rate = 25%), 120 of which were used as critical switch trials. At a critical switch, the stimuli on the current (switch) and the preceding trial were both noncognates. In this way we could look at “purer” switches because the language borders are less clear for cognates. Within a list, each noncognate item occurred six times on a critical switch, three times in each language. We pseudo-randomized all the items in each block using the program MIX [48], with the following requirements: (1) there were no more than four subsequent trials with the same cognate status; (2) subsequent trials are semantically, phonologically, and pragmatically unrelated; (3) repetition of a picture was separated by at least four intervening trials; (4) there were no more than six subsequent trials in the same language; (6) there were no subsequent switch trials. A second list was constructed by reversing the block order of the first list. Based on the pseudo-randomized lists, we further selected 264 critical repeat trials in each list by (1) excluding post-switch trials; (2) matching the number of cognate and noncognate trials between languages; (3) matching the ordinal position of cognate and noncognate trials within sequences of repeat trials of each language (all *ps* > .129). Within the 264 critical switch trials per list, each of the 80 items occurs minimal once and maximally five times after randomization.

### Procedure

We seated the participants in a sound-proof booth and ran the experiment using the software package *Presentation* (Version 17.0, Neurobehavioural System Inc, Berkeley, U.S.).
The computer screen (Benq XL2420Z, 24-inch screen) was set to grey, with a resolution of 1920 × 1080 pixels, at a refresh rate of 120 Hz. Each session consisted of four parts: item familiarization, cue familiarization, speed training with time pressure, and experimental blocks. During item familiarization, participants saw each picture and named it in Dutch (block 1) or English (block 2); if they were unable to name it, they were told the correct answer and asked to remember it and name it again. Cue familiarization served the training of the color-language association; and in the speed training, we introduced time pressure with the aim that participants would make more speech errors. For a more detailed description of the procedure for familiarization and speed training, we refer to Zheng et al. [34].

We recorded participants’ EEG during the experimental blocks. Each trial started with the 250 ms presentation of a fixation cross, followed by a blank screen with a jitter of 250-500 ms. Then, the picture appeared in the center of the screen, with a 100-pixel-wide frame around the picture whose color represented the response language (i.e., red and yellow indicated Dutch, and green and blue indicated English, or vice versa). Two colors were used to cue each language to avoid a confound of language switch and color switch [49]. We counterbalanced the assignment of colors to the response language across participants. The picture and the frame stayed on the screen until 550 ms after the voice key (Shure SM-57 microphone) had registered the onset of speech. If the voice key was not triggered within 2000 ms, the stimulus stayed on the screen for a total of 2550 ms. After another jittered blank screen of 250-500 ms, the next trial began. After each block, participants received feedback on their performance (e.g., speed). We instructed them to name the pictures as quickly as possible in the language indicated by the cue, and also not to correct themselves when they said something wrong. All the instructions were in English.

After the EEG measurement, participants completed the LexTALE vocabulary test in English and a language background questionnaire. The entire session took approximately 2.5 hrs.

### EEG recording

We recorded EEG using an elastic cap containing 57 active Ag-AgCl electrodes based on the international 10-20 system (ActiCAP 64Ch Standard-2, Brain Products). Seven additional electrodes were placed on both mastoids (reference), the forehead (ground), next to the eyes (EOG), and next to the upper lip and the throat (EMG). EEG signals were referenced to the left mastoid electrode online and re-referenced to the average of the right and left mastoid electrodes offline. EOG was measured with the electrodes placed above and below the right eye (to monitor for vertical eye movements) and on the left and right temples (to monitor for horizontal eye movements). EEG, EOG and EMG signals were sampled at a frequency of 500 Hz and online-filtered with a low cut-off of 0.016 Hz and a high cut-off of 125 Hz. Impedances for EEG electrodes were kept below 20 kΩ.

### EEG preprocessing

We performed all EEG analyses using the Fieldtrip open source Matlab toolbox [50] and custom analysis scripts using Matlab v.8.6.0 (R2015b, The Math Works, Inc).

As mentioned before, the potential speech movement related artifacts during response-locked EEG has been considered an issue for ERP analysis in language production [19]. Therefore, we performed a pilot study (5 participants) to evaluate three different methods for removing muscle artifacts associated with overt speech production: canonical correlation analysis (CCA), independent component analysis (ICA), and low-pass filtering (high cut-off = 10 Hz). We measured participants’ EEG during a Stroop task which required either manual (block 1) or vocal responses (block 2). After applying one of the three methods, or none of them, to the EEG data of the vocal Stroop task, we compared its response-locked ERPs with those from the manual Stroop task. To our surprise, all the three methods turned out to be unnecessary regarding our data: the uncorrected data of the vocal task that were submitted to only a standard artifact rejection procedure (see below) gave a clear ERN that was comparable to that in the manual task. For example, the muscle artifacts seemed to be too small to be detected by the CCA algorithm. The same analyses were performed to our data in the main experiment and led to the same conclusion. Therefore, we decided to simply apply an extra round of visual inspection to remove trials with muscle artifacts (see below).

The EEG signal was preprocessed as follows: First, we segmented the continuous EEG into stimulus-locked epochs from 200 ms before to 2500 ms after each picture onset. The data were then re-referenced and band-pass filtered with a low cut-off of 0.1 Hz and a high cut-off of 30 Hz. Trials with atypical artifacts (e.g., jumps and drifts) were rejected after visual inspection; EOG artifacts (eye blinks and saccades) were removed using ICA. After that, we applied another round of visual inspection to remove trials with remaining artifacts (e.g., muscle artifacts due to articulation). Baseline correction was applied based on the average EEG activity in the 200 ms interval before picture onset and the data were further segmented into stimulus-locked epochs (from 200 ms before to 500 ms after each picture onset) and response-locked epochs (from 500 ms before to 500 ms after each vocal response, see below for the offline adjustment of speech onset). Individual EEG channels with bad signals were disabled before ICA for EOG artifacts and interpolated by a weighted average of the data from neighboring channels of the same participant. On average, we discarded 7.3% of the stimulus-locked data, 6.4% of the response-locked data, and 1.6 channels per participant. Eleven channels (AF8, F7, FT7, FT8, CP5, T8, TP8, P5, P7, P8, and PO7) that were repaired in more than one participant were excluded from the group-level analyses.

We averaged all the stimulus-locked and response-locked segments for each condition and each participant, respectively. Participants with less than 15 remaining trials in any condition were excluded from the EEG analysis.

### Statistical analysis

For the behavioral data, we used error rates and RTs as dependent variables. Participants’ responses were coded as correct, fluent responses, and incorrect responses. Incorrect responses were further categorized into language selection errors (i.e., complete, fluent responses in the nontarget language) and another twelve types of errors, such as self-corrections, disfluencies, or using a wrong word in the correct language. We re-measured speech onset manually in Praat [51] and discarded RT outliers based on individual participants’ performance, aggregated by language and trial type (switch vs. repeat). Correctly responded trials with a RT deviating more than three standard deviations from the respective participants’ condition mean were defined as another type of error (i.e., RT outliers, see S1 Appendix B for all the categories and the percentages of each type of error). In the error analysis, we excluded trials that could not be classified as either switch or repeat (trials at the beginning of each block and trials following language selection errors or other interlingual errors; see S1 Appendix B). In the RT analysis, we excluded all error trials and post-error trials.

The statistical analysis of the behavioral data was carried out using mixed-effects models with the lme4 package (Version 1.1.13) [52] in R (Version 3.4.1) [53]. For the repeat trials, we sum-coded the factors language (L1 vs. L2) and cognate status (cognate vs. noncognate) and included them as fixed effects in the models. For the switch trials, only language was included as a fixed effect. Participants and items were included as random effects in both analyses. We started all the analyses with a maximal random-effects structure – that is, models including random intercepts and random slopes for all fixed effects and their interactions for both participants and items [54]. Only when the model with the maximal random-effects structure did not converge, we simplified it by first removing the interactions and if necessary the main effects in the random structure (see S1 Appendix C for the final models used for analyses). We used generalized linear mixed models (GLMEMs) to analyze error rates as well as RTs. Compared to linear mixed effects models, GLMEMs can account for the right-skewed shape of the RT distribution without the need to transform and standardize the raw data [55]. For each analysis, we reported Wald’s *z*-scores, *t*-scores and their associated *p*-values.

The statistical analysis of the ERP data was run using a nonparametric cluster-based permutation test [56]. The method controls for the false alarm rate when a large number of comparisons have to be made to evaluate the ERP data at multiple channels and multiple time points. On the critical repeat trials, we compared correct responses of cognate vs. noncognate naming; on the critical switch trials, we compared language selection errors vs. correct responses. Post-error trials were excluded in both analyses. We try to briefly describe the procedure of the cluster-based permutation test here, but refer to the original article [56] for details.

We first compared the two conditions with a paired-samples t-test (two-tailed) at each spatiotemporal sample (i.e., per channel per time point). Then, we selected all samples whose *p*-values were smaller than the given threshold of .05. Afterwards, those selected samples which were spatiotemporally adjacent were grouped as clusters. For each cluster, the t-values of all the samples were summed, yielding the cluster-level statistic. Then the cluster with the maximum cluster-level statistic was selected to compare against a permutation distribution. The permutation distribution is constructed through random partitioning the original data for 1000 times and determining spatiotemporal clusters with their cluster-level statistic with the same procedure as described above. For the selected cluster, its *p*-values were calculated as the proportion of random partitions (out of 1000) that yielded a larger cluster-level statistic than its statistic. We consider *p*-values below .05 (two-tailed) significant.

We focused on two ERP components which are taken to reflect the processing of error or conflict: the stimulus-locked N2 and the response-locked ERN. For the analyses of the ERN, we applied statistical tests to three fronto-central channels at which the ERN is typically reported (Fz, FCz, and Cz). In the action literature, the ERN is mostly reported in the time window of 0 to 100 ms post response onset. However, due to the complex nature of speech production, the onset of the ERN might have a different timing relative to the speech onset. Since the previous literature on language production [11, 16] did not give a clear indication of the onset of the ERN, we took a more conservative approach and applied the permutation test to the full time window (i.e., 500 ms pre to 500 ms post speech onset). For the analysis of the N2, we have more consistent information about its time window from the literature (i.e., peaking around 200 ms post stimulus, e.g., [37–38]). Therefore, we limited the analysis to a narrower time window (i.e., 150 ms to 350 ms post stimulus onset), but applied it to all the available electrodes given the widely distributed topography of the different N2s (i.e., fronto-central N2 and posterior N2).

## Behavioral results

### Analysis of switch trials

Speakers made different types of speech errors on 21.4% of all trials (i.e., including both switch and repeat trials), including responses in the nontarget language (e.g., say the Dutch word “boom” instead of the English word “tree”; language selection errors) on 16.4% of the trials. On critical switch trials, language selection errors reached an average rate of 37.3%.

In general, speakers made more language selection errors when they had to switch to their L1, Dutch (*M*_L1_ = 43.5%, *SD*_L1_ = 15.6%) than when switching to their L2, English (*M*_L2_ = 30.0%, *SD*_L2_ = 16.1%; β = 0.33, *SE* = 0.10, *z* = 3.23, *p* = .001). They were also slower when switching from the L2 to the L1 (*M*_L1_ = 825 ms, *SD*_L1_ = 123 ms) than vice versa (*M*_L2_ = 767 ms, *SD*_L2_ = 96 ms; β = 30.66, *SE* = 11.54, *t* = 2.66, *p* = .008).

### Analysis of repeat trials

Fig 1 shows the violin plots for the language selection error rates and the RTs on the repeat trials. Table 3 gives the statistics from the GLMEMs for the language selection error rates and the RTs on the repeat trials.

**Fig 1.**
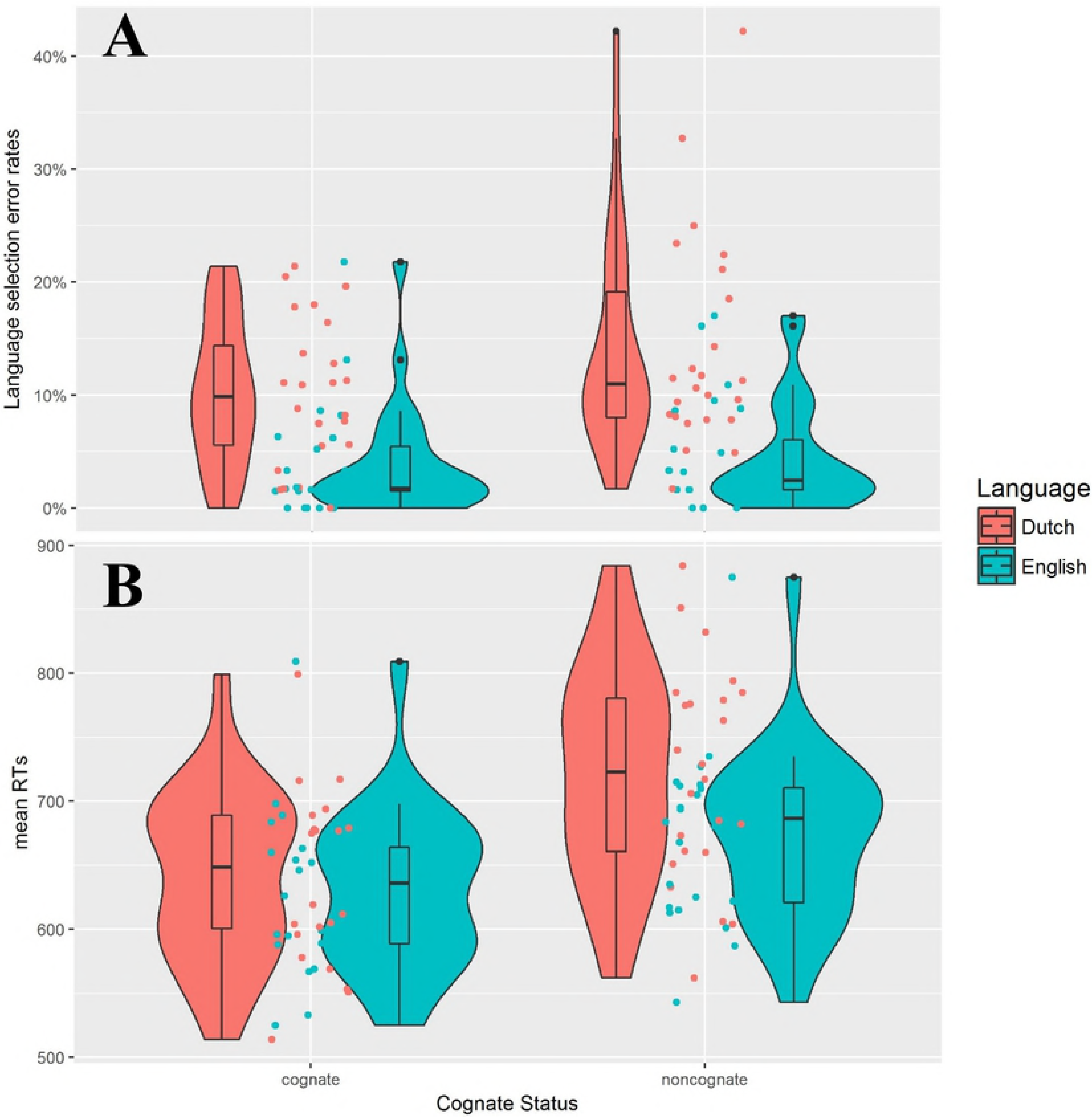
Violin plots with individual data distributions for language selection error rate and mean RT. (A) Language selection error rate and (B) mean RT (in ms), grouped by language (Dutch vs. English) and cognate status (cognate vs. noncognate). The outer shapes represent the distribution of individual data, the thick horizontal line inside the box indicates the median, and the bottom and top of the box indicate the first and third quartiles of each condition.

**Table 3.** Statistics from the GLMEMs for the Language Selection Error Rate (%) and the Reaction Time (ms) on Repeat Trials.

| | | *Mean (SD)* | | $\beta$ | *SE* | *z- or t-value* | *p* |
| --- | --- | --- | --- | --- | --- | --- | --- |
| ER | Language | L1 | 12.1 (8.3) | **0.67** | **0.14** | **4.84** | **<.001** |
|  |  | L2 | 4.9 (4.9) |  |  |  |  |
|  | CS | cognate | 7.1 (6.5) | -0.14 | 0.10 | -1.46 | .144 |
|  |  | noncognate | 9.3 (8.9) |  |  |  |  |
| | Language × CS | | | -1.02 | 0.10 | -0.26 | .799 |
| RT | Language | L1 | 682 (85) | **23.48** | **5.21** | **4.51** | **<.001** |
|  |  | L2 | 652 (68) |  |  |  |  |
|  | CS | cognate | 636 (64) | **-35.27** | **4.88** | **-7.23** | **<.001** |
|  |  | noncognate | 698 (79) |  |  |  |  |
| | Language × CS | | | **-10.06** | **4.95** | **-2.03** | **.042** |

*Note.* CS = cognate status. ER = error rate. RT = reaction time.
Significant effects are highlighted in bold.

On repeat trials, speakers also made more language selection errors when naming in the L1 than in the L2. However, error rates were comparable when naming cognate and noncognate words. There was no interaction between language and cognate status.

As for RTs, speakers were also slower when naming in the L1 than in the L2. Contrary to the language selection error rates, speakers were actually faster in cognate naming than noncognate naming. There was also a significant interaction between cognate status and language. A follow-up analysis for each language showed that the cognate facilitation effect was larger in the L1 (*M*_L1cog_ = 642 ms, *SD*_L1cog_ = 67 ms; *M*_L1noncog_ = 722 ms, *SD*_L1noncog_ = 83 ms; β = −45.08, *SE* = 9.68, *t* = −4.66, *p* < .001) than in the L2 (*M*_L2cog_ = 632 ms, *SD*_L2cog_ = 63 ms; *M*_L2noncog_ = 673 ms, *SD*_L2noncog_ = 67 ms; β = −27.49, *SE* = 6.46, *t* = −4.25, *p* < .001).

## EEG results

### Response-locked analysis

#### Analysis of switch trials

On critical switch trials, we compared language selection errors with correct responses. Fig 2 shows the response-locked ERPs and topographies for both conditions.

**Fig 2.**
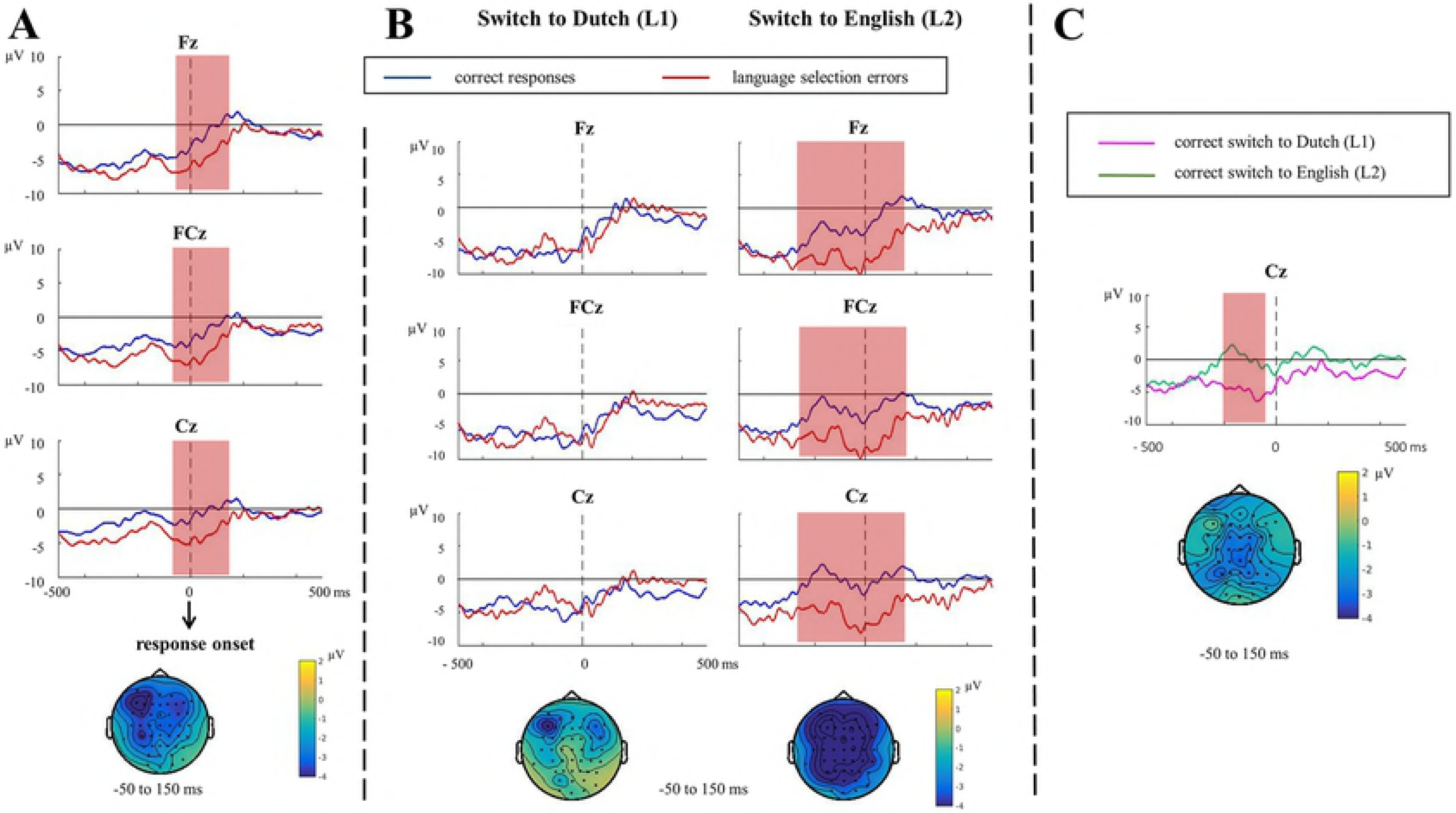
Response-locked ERPs and topographies for switch trials. (A) Response-locked ERPs and topographies for language selection errors vs. correct responses (*N* = 24). (B) Response-locked ERPs and topographies for language selection errors vs. correct responses when switching to L1, Dutch and switching to L2, English (*N* = 21). (C) Response-locked ERPs and topographies for correct responses when switching to L1, Dutch vs. switching to L2, English (*N* = 21). When the ERN/CRN effect was significant between conditions, the time windows associated with the statistically significant effect are marked in light red. Topographies of the difference between the two conditions are presented for each contrast. To better compare the topographies between contrasts, we used the same time window to which the ERN effect was associated in A also for B and C. Baseline correction was applied based on the 200 ms interval before picture onset (not shown in the current figure).

For the response-locked data, the cluster-based permutation test revealed a significant difference between language selection errors and correct responses (*p* = .004), with the difference being most pronounced around 50 ms pre-to 150 ms post-response onset (the ERN effect, Fig 2A).

To test whether the ERN effect was moderated by switching directions (i.e., switching to L1 vs. switching to L2), we split the data by languages and applied the same tests separately (Fig 2B). Given the limited number of remaining trials per language, we accepted a minimum of six trials per cell [57] and thus had 21 participants’ data available for the analysis. Results showed a significant difference between language selection errors and correct responses when switching to the L2, English (*p* = .006), with the effect being even more wide-spread in time (250 ms pre-to 150 ms post-response onset). In contrast, no ERN effect in switching to the L1, Dutch, was found in the response-locked analysis (*p* = .516). Therefore, when making a language selection error in switching to the L2, speakers showed a larger ERN effect than when switching to the L1 (*p* = .018). Moreover, visual inspection suggested a possible difference between the correct responses in the L1 and the L2. Therefore, we applied the same cluster-based analysis to the correct responses between the two switching directions (Fig 2C) and found a difference between the two conditions (*p* = .010): When speakers correctly switched to the L1, their response-locked ERPs were more negative (i.e., larger CRN) than when they correctly switched to the L2. The effect was most pronounced around from 200 to 50 ms pre-response onset.

However, the EEG data during overt speech can be noisier than Pontifex et al. [57] and thus may affect our conclusions about the ERP effects. Therefore, we verified the current analysis with a minimum of 10 trials per participants per condition and 13 available participants. All the main results about the ERN/CRN persisted. Also, for the sake of consistency, we applied the same behavioral analysis again to this subset of 21 participants. The pattern of results did not differ from the one in the full dataset.

#### Analysis of repeat trials

On critical repeat trials, we compared correct responses of cognate vs. noncognate naming. Fig 3 shows the response-locked ERPs and topographies for both conditions.

**Fig 3.**
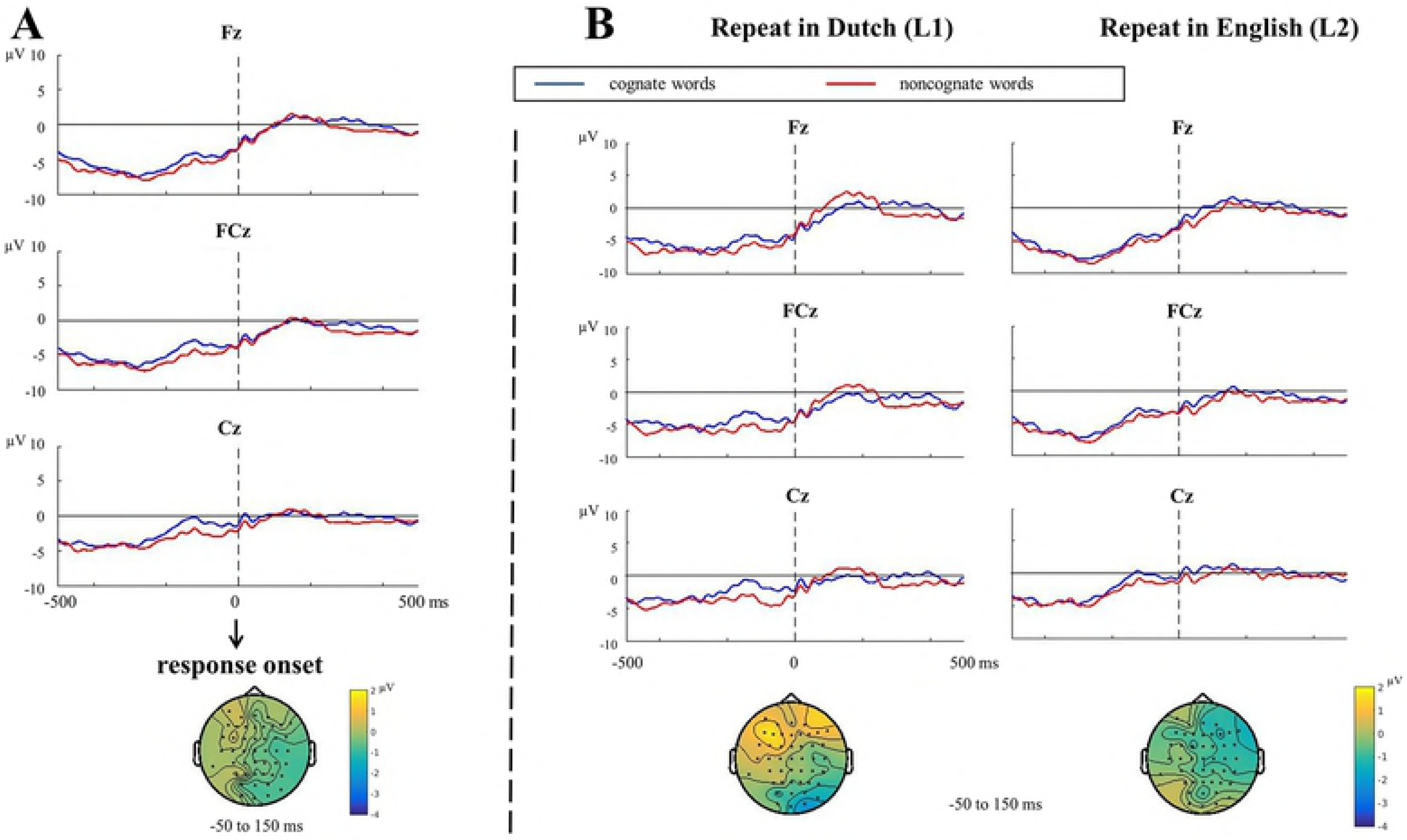
Response-locked ERPs and topographies for repeat trials. (A) Response-locked ERPs and topographies for correct naming of cognates vs. noncognates (*N* = 24). (B) Response-locked ERPs and topographies for cognate vs. noncognate naming when repeating in L1, Dutch and in L2, English (*N* = 24). Topographies of the difference between the two conditions are presented for each contrast. For better comparison, we used the same time window to which the ERN effect was associated in the analysis of switch trials. Baseline correction was applied based on the 200 ms interval before picture onset (not shown in the current figure).

No significant difference between correct responses to cognate and noncognate words was found in the response-locked data (*p* = .078, Fig 3A). We also compared the potential CRN effect for cognates vs. noncognates between languages (repeat in L1 vs. repeat in L2, Fig 3B). There was no significant CRN effect either in English (*p* = .510) or in Dutch (*p* = .080), and there was also no difference between languages (*p* = .138).

### Stimulus-locked analysis

#### Analysis of switch trials

Fig 4 shows the stimulus-locked ERPs and topographies for language selection errors vs. correct responses on critical switch trials.

**Fig 4.**
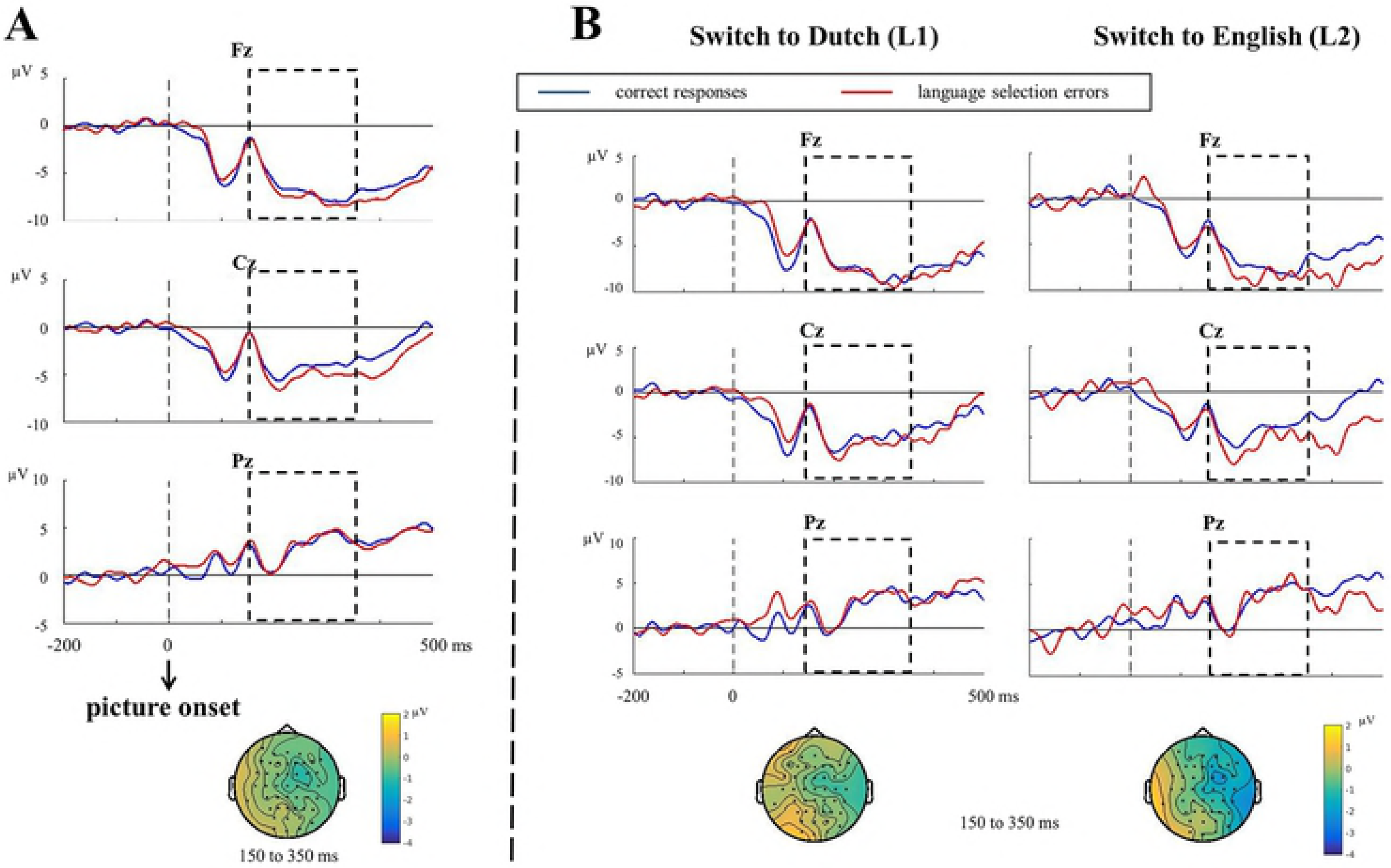
Stimulus-locked ERPs and topographies for switch trials. (A) Stimulus-locked ERPs and topographies for language selection errors vs. correct responses (*N* = 24). (B) Stimulus-locked ERPs and topographies for language selection errors vs. correct responses when switching to L1, Dutch and switching to L2, English (*N* = 21). The time window used for testing the N2 effect (150 to 350 ms) is marked by an empty frame. Topographies of the difference between the two conditions are presented for each contrast.

Cluster-based permutation tests applied to the stimulus-locked data revealed no N2 effect between language selection errors and correct responses (*p* = .573, Fig 4A). We also compared stimulus-locked data between the two switching directions (Fig 4B). Results showed no N2 effect either in switching to the L1 (*p* = .655) or to the L2 (*p* = .438). There was also no difference between the two switching directions (*p* = .488).

We verified the analysis again with a minimum of 10 trials per participants per condition and 13 available participants: Now, an N2 effect was found in language selection errors compared to correct responses when switching to the L2 (*p* = .018), mostly pronounced between 290 to 350 ms post stimulus onset, central electrodes. The difference between switching directions, however, was not significant.

#### Analysis of repeat trials

Fig 5 shows the stimulus-locked ERPs and topographies for correct responses of cognate vs. noncognate naming on critical repeat trials.

**Fig 5.**
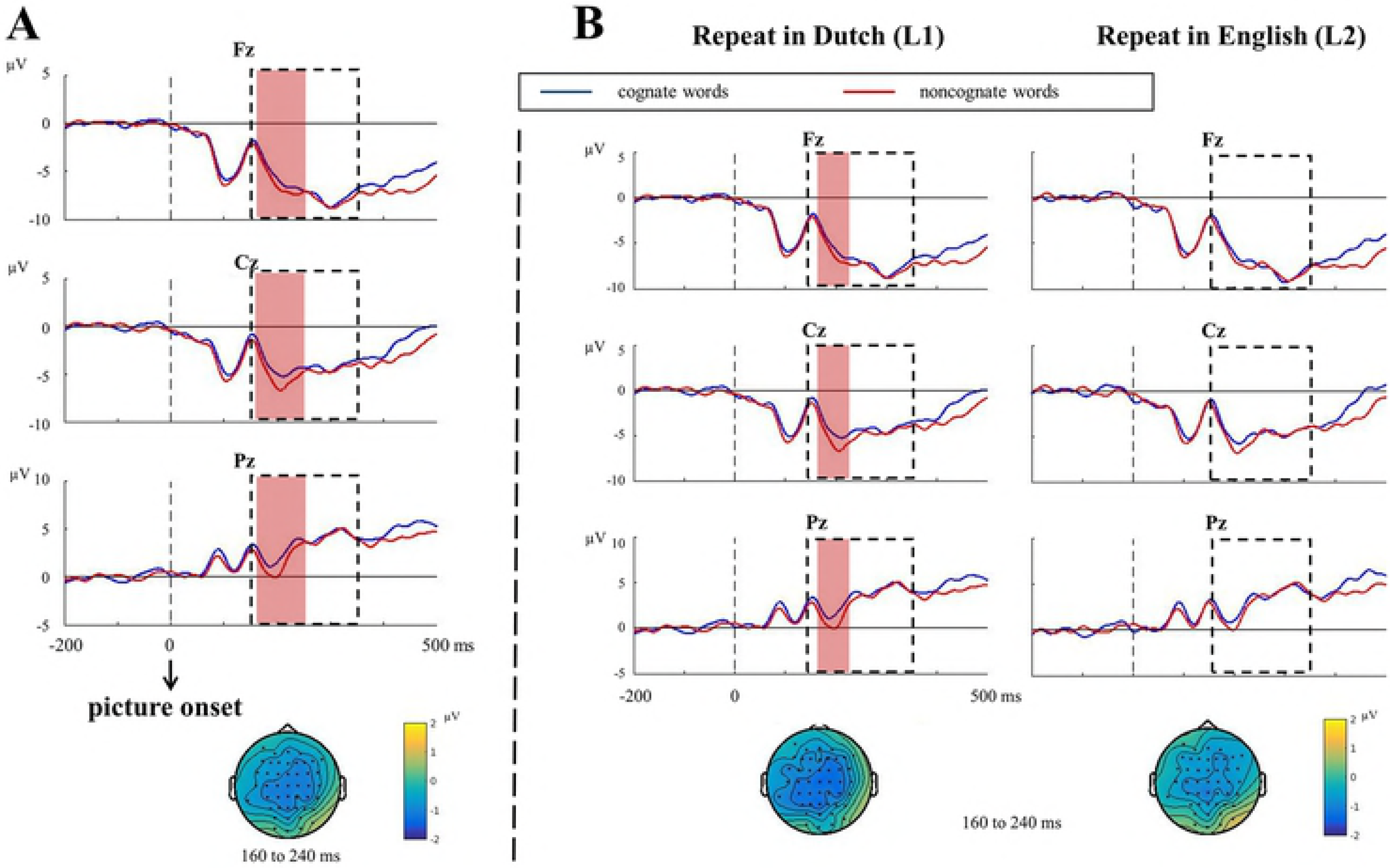
Stimulus-locked ERPs and topographies for repeat trials. (A) Stimulus-locked ERPs and topographies for correct responses of cognate vs. noncognate naming (*N* = 24). (B) Stimulus-locked ERPs and topographies for cognate vs. noncognate naming when repeating in L1, Dutch and in L2, English (*N* = 24). The time window used for testing the N2 effect (150 to 350 ms) is marked by an empty frame. When the N2 effect was significant between conditions, the time windows associated with the statistically significant effect are marked in light red. Topographies of the difference between the two conditions are presented for each contrast. To better compare the topographies between contrasts, we used the same time window to which the N2 effect was associated in A.

The stimulus-locked analysis showed an N2 effect in noncognate words compared to cognate words (*p* = .006), which was most pronounced between 160 to 240 ms post stimulus onset at central electrodes (Fig 5A). When comparing between languages (Fig 5B), an N2 effect was revealed for noncognate words compared to cognate words in L1, Dutch (*p* = .018) that was most pronounced between 180 to 210 ms post stimulus onset at central electrodes, but no difference was revealed between cognate and noncognate words in L2, English (*p* = .621). The difference between languages, however, was not significant (*p* = .849).

## Discussion

In the current study, we investigated how bilingual speakers monitor their speech errors and control their languages in use. We found that bilingual speakers were slower and made more language selection errors when switching from the L2 to the L1 than vice versa, replicating previous findings on the reversed dominance effect [30–32, 34, 37]. This is probably because when speaking in the weaker L2, more cognitive control is needed (e.g., to inhibit the nontarget L1 and/or to enhance the target L2) than speaking in the stronger L1. Therefore, when switching back to the L1, it is more difficult to overcome the residual control [34].

We also observed a robust ERN effect after language selection errors compared to correct responses. Compared to previous research on action monitoring [7–8, 24], the ERN effect in the current study was observed in a rather early time window (−50 to 150 ms relative to response onset), suggesting that speech monitoring takes place earlier than actual articulation (i.e., during speech planning).

To test the conflict-based monitoring model [21], we compared the ERN effect between switching to L2 and switching to L1. According to the conflict-based model, higher error rates and longer RTs suggest more response conflict when switching from L2 to L1 than vice versa, and therefore, a larger ERN effect should be expected when incorrectly switching to L1 (i.e., the high-conflict condition) than to L2 (the low-conflict condition). However, we observed the opposite effect, namely, a larger ERN effect following language selection errors in switching to L2 than to L1 (also opposite to the behavioral error effect). Therefore, our result challenges the conflict-based model [21] (but see [24]). We will discuss this in more depth below. In line with our behavioral finding that switching from L2 to L1 is more difficult than in the opposite direction (i.e., longer RTs), we also observed a larger CRN, an equivalent of the ERN on correct trials, when speakers switched correctly from L2 to L1 than vice versa. This reflects a greater general difficulty in correctly switching from L2 to L1 than vice versa (see also [13]). As for the stimulus-locked analysis, we did not find an N2 effect in language selection errors compared to correct responses. There was also no N2 effect in the between-language analysis. This suggests a possible dissociation between the ERN and the N2 in error monitoring (but not necessarily in conflict monitoring, see [24]).

We also tested the conflict-based model by comparing the CRN difference between correct responses for cognate and noncognate naming. According to the predictions of the model, noncognates should give rise to a larger CRN because more conflict is involved when naming them compared to form-overlapping cognates. However, we did not find any difference in the CRN between the two conditions. There was also no difference in error rates between cognates and noncognates. Yet, in line with the expectations, we did observe faster naming for cognate words than noncognate words, replicating the cognate facilitation effect [29, 40–41]. This cognate facilitation effect was larger in L1 than in L2 in a mixed language context, suggesting that the dominance of the L1 is reversed in this task and L1 is thus more likely to be influenced by L2 than vice versa (see also [30]). We also observed an N2 effect in the correct naming of noncognate compared to cognate words. The effect was restricted to the L1. Our ERP results are opposite to the results obtained by Christoffels, Firk, and Schiller [30], who observed a larger N2 in cognate compared to noncognate naming.

Although the conflict-based monitoring model [21] is challenged by our finding of a larger ERN/CRN effect in the more accurate switching condition (L1 to L2), the original conflict monitoring theory [24] did not associate a larger ERN effect with the condition in which errors are particularly likely (e.g., switching from L2 to L1). This is because the original conflict-monitoring theory differs from the conflict-based model of speech monitoring in terms of the exact point in time when the conflict associated with the ERN is assumed to be detected. According to the original conflict-monitoring theory [24], the ERN is the result of post-error conflict between the actually committed incorrect response and the intended correct response. Therefore, when a correct response is more likely but an error is nevertheless committed, the ERN amplitude is increased by the higher activation built up in favor of the correct response following an actual error (i.e., more post-error conflict). For example, the conflict monitoring theory predicts less post-error conflict, and thus a smaller ERN, on (high-conflict) incongruent trials than on (low-conflict) congruent trials in a Flanker task (see also [58]). While the original theory makes predictions based on post-response conflict, the conflict-based model of monitoring in language production [21] assumes conflict to take place during the planning stage of the word (i.e., pre-response conflict), in particular, between words during word selection and between phonemes during phoneme selection. This leads to their opposing predictions about the ERN amplitude in terms of conflict.

Whereas the conflict-based model of monitoring in language production [21] is challenged by our data, previous ERN studies of language production challenge the original conflict-monitoring theory. Ganushchak and Schiller [14] reported a larger ERN on errors following semantically related (i.e., more conflict) compared to unrelated distractors in a picture-word interference task. Using a semantic blocking paradigm, the same authors also found a larger ERN in semantically related blocks compared to unrelated blocks [11], suggesting that the amplitude of the ERN can be proportional to the amount of pre-response conflict.

Our current results on the ERN are in conflict with these previous results on the relation of ERN and error rates / conflict, but in line with theories of how the monitoring system predicts errors and uses such information for reinforcement learning. To optimize performance, the monitoring system learns to predict errors in ongoing events [59] and adjusts its prediction for further learning [60]. In a given context, the prediction of errors is made based on context features such as error likelihood [59] (but see [61]). The monitoring system gets altered when errors are more likely (as reflected by a larger CRN). When an error occurs without being predicted, reinforcement learning occurs and such information is used to refine ongoing predictions (as reflected by a larger ERN). Applied to the language switching scenario, switching from L2 to L1 is more difficult than switching in the opposite direction, and thus more likely to cause a language selection error. In order to switch properly, the monitoring system enhances its activity to match the predicted demand. Early warning signals (as reflected by a larger CRN) are sent for recruiting and regulating cognitive control and monitoring [59]. This is consistent with our finding of a larger CRN during correctly switching to L1 than during switching to L2. When a language selection error is actually committed as predicted when switching from L2 to L1, less adjustment to the monitor prediction is necessary and thus the ERN effect (i.e., the difference between actual errors and correct responses) is smaller/absent. On the other hand, when switching from L1 to L2, it is relatively easier to make a switch and fewer errors are likely, thus the monitor activity remains low (as reflected by a smaller/absent CRN). Nevertheless, when an error is indeed committed, such an unexpected action will be used for reinforcement learning in order to improve future performance [60]. This is reflected by a larger ERN following an error when switching from L1 to L2 than vice versa.

In summary, as a successful first attempt to investigate the error monitoring process in bilingual switching, the current study found a robust ERN effect for language selection errors. Moreover, we found that the ERN effect is larger for a language selection error when switching to L2 (i.e., low-conflict condition) than switching to L1 (i.e., high-conflict condition), which challenges the conflict-based monitoring model [21]. Rather, the data suggest a role of the ERN in error prediction and reinforcement learning not only in the monitoring of actions, but also in bilingual language control.

## Acknowledgement

The authors would like to thank Robert Hartsuiker, Vitória Piai, Eric Maris and Jana Klaus for their helpful discussions during the project.

## Supporting information

**S1 Appendix A. Materials**. A full list of cognate and noncognate words used in the study.

**S1 Appendix B. Error coding**. All the categories and the percentages of each type of speech errors coded in the study.

**S1 Appendix C. Linear mixed effect models**. The linear mixed effect models used for analyses in the study.

